# Improving emotional-action control by targeting long-range phase-amplitude neuronal coupling

**DOI:** 10.1101/2020.06.04.129569

**Authors:** Bob Bramson, Hanneke den Ouden, Ivan Toni, Karin Roelofs

## Abstract

Control over emotional action tendencies is essential for every-day interactions. This cognitive function can fail during socially challenging situations, and is chronically attenuated in social psychopathologies such as social anxiety and aggression. Previous studies have shown that control over social-emotional action tendencies depends on phase-amplitude coupling between prefrontal theta-band (6 Hz) rhythmic activity and broadband gamma-band activity in sensorimotor areas. Here, we delivered dual-site phase-coupled brain stimulation to facilitate theta-gamma phase-amplitude coupling between frontal regions known to implement that form of control, while participants were challenged to control their automatic action tendencies in a social-emotional approach/avoidance-task. Participants had increased control over their emotional action tendencies, depending on the relative phase and dose of the intervention. Concurrently measured fMRI effects of task and stimulation, and estimated changes in effective connectivity, indicated that the intervention improved control by increasing the efficacy of anterior prefrontal inhibition over sensorimotor cortex. This enhancement of emotional action control provides causal evidence for a phase-amplitude coupling mechanism guiding action selection during emotional-action control. More generally, the finding illustrates the potential of physiologically-grounded interventions aimed at reducing neural noise in cerebral circuits where communication relies on phase-amplitude coupling.

## Main text

The ability to control emotional actions is paramount for successful engagement in human social interactions (*1*). Long-standing theorising and accumulating empirical evidence indicate that affective cues automatically activate approach-avoidance action tendencies (*2, 3*). Effective emotion control requires the cognitive capacity to suppress those automatic action tendencies and to select an alternative course of action (*4, 5*). The importance of emotional-action control becomes apparent when it is disrupted: In social psychopathologies such as social anxiety, the inability to override social avoidance tendencies constitutes the core maintaining factor of the disorder (*6*). There is great interest in potentiating this cognitive capacity to enhance treatment efficacy, as well as to help professionals exposed to socially challenging situations. However, improving human emotional-action control has proven difficult (*7*). In this study, we use a brain stimulation intervention designed to enhance synchrony within a cerebral circuit known to support emotional-action control (*8, 9*). By modelling how effective and structural connectivity of that cerebral circuit mediates the behavioural effects of the intervention, we provide an account of its neural effects, paving the way for physiologically-grounded therapeutic interventions in social-emotional disorders (*10*).

Previous non-invasive brain stimulation interventions have been successful in *reducing* emotional-action control. This was achieved by disrupting neural activity – putatively by injecting neural noise (*11*) – in a region known to coordinate emotional-action control: the anterior prefrontal cortex (aPFC) (*9*). Here, we explore whether it is possible to *enhance* emotional-action control by using brain stimulation aimed at reducing neural noise, targeting a cortical circuit known to regulate those action-tendencies (*7–9*). This gain-of-function intervention is grounded on recent insights showing that emotional-action control requires neural synchronisation between aPFC theta-band rhythm and sensorimotor broadband gamma activity (*8, 12, 13*). We reasoned that endogenous neural synchronization might be enhanced by applying separate time-varying electric fields (transcranial alternating current stimulation; tACS (*14*)) to aPFC and sensorimotor cortex (SMC). tACS influences spike timing of individual neurons (*14, 15*), and entrains neural rhythms to its frequency and phase (*16, 17*). We applied dual-site phase-coupled tACS to enhance the endogenous synchronization of SMC gamma-band power (75 Hz) with the peaks of aPFC theta-band rhythm (6 Hz) evoked during emotional-action control (*8, 18, 19*). Enhanced phase-amplitude coupling would reduce neural noise in aPFC-SMC communication (*10, 13*), allowing for improved control over emotional-action tendencies. Crucially, we apply this aPFC-SMC stimulation while 41 participants perform an emotional-action control task (Figure 1A). We contrast the online behavioural and neural effects of in-phase, anti-phase, and sham couplings between the power envelope of SMC gamma stimulation and the peaks of aPFC theta stimulation (Figure 2A;B). Importantly, concurrent whole-brain BOLD-fMRI quantified local and remote dose-dependent cerebral effects of the electrical stimulations (*20*). tACS effects on each participants’ cerebral connectivity were further qualified using dynamic causal modelling (*21*), informed by MR-tractography (*22*).

**Figure 1:**
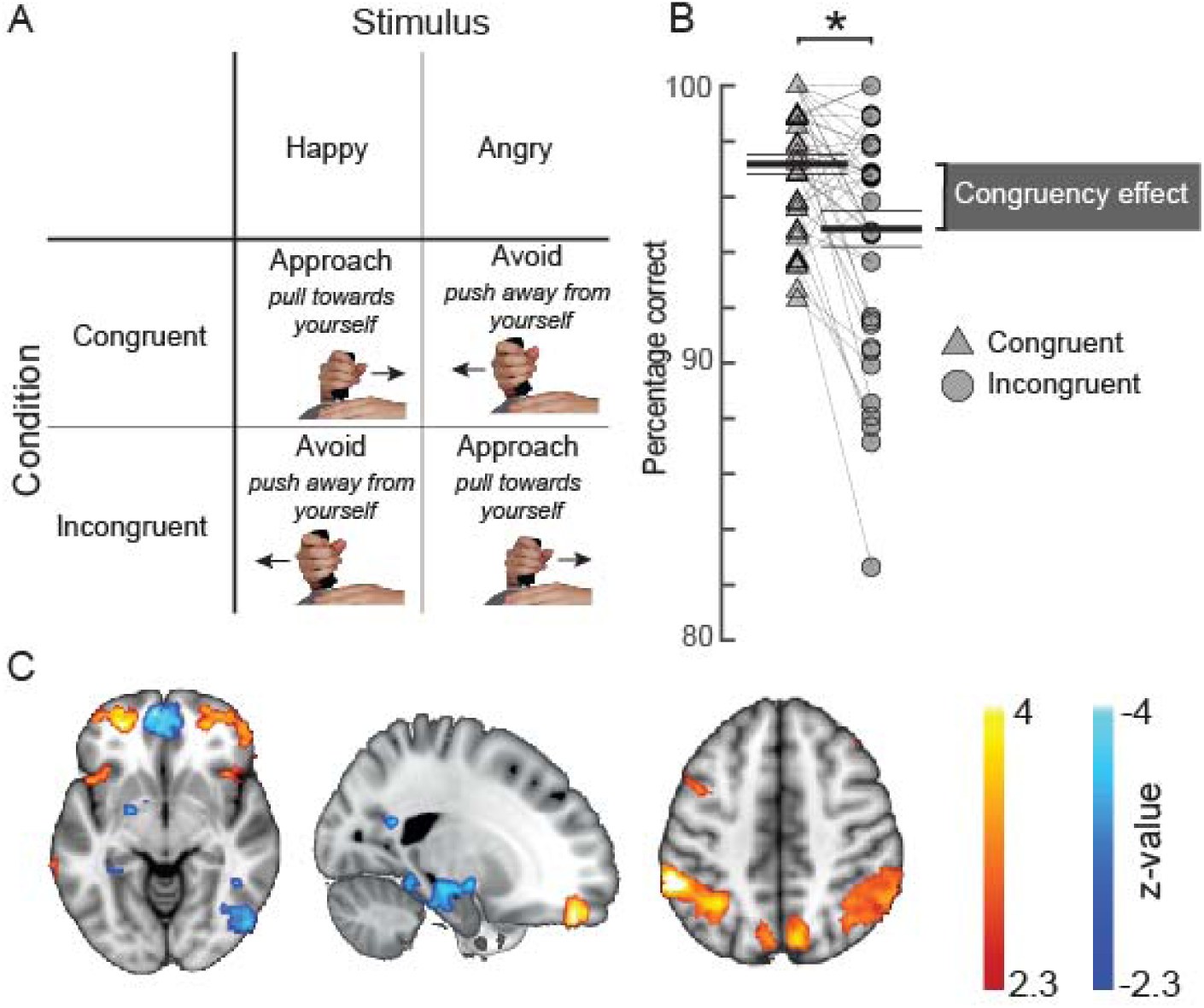
Behavioural and cerebral effects of the approach-avoidance task used to manipulate control over emotional action tendencies. A: Conceptual visualisation of the approach-avoidance task. Participants pushed- or pulled a joystick away- or towards themselves to approach- or avoid happy and angry faces. Approaching angry- and avoiding happy faces is incongruent with action tendencies to approach appetitive and avoid aversive situations. B) Behavioural results in the sham condition of the task. Participants make more errors in the incongruent trials (circles) than in the congruent trials (triangles). Black lines visualise the mean and standard error of the mean. Grey bar depicts the congruency effect. C) Approach-avoidance congruency-related BOLD changes (p<.01 cluster-level inferences corrected for multiple comparisons). Trials involving responses incongruent with automatic action tendencies showed stronger BOLD signal in anterior prefrontal- and parietal areas, and reduced signal in left amygdala/hippocampus and medial prefrontal cortex.

**Figure 2:**
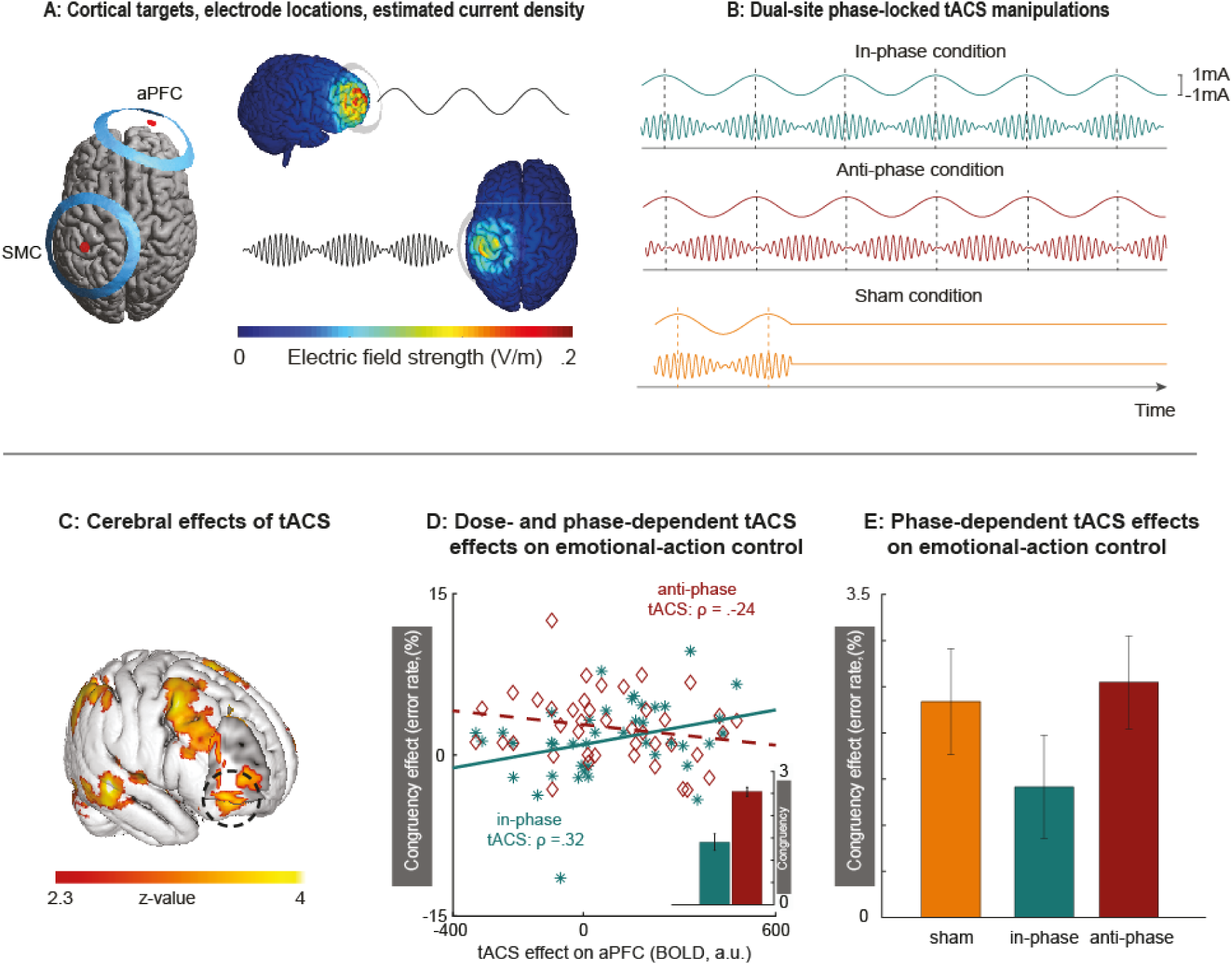
Behavioural effects of dual-site phase-coupled tACS on emotional-action control are dose- and phase-dependent. A) Two sets of ring electrodes were placed over right aPFC and left SMC. Modelling of current density showed that stimulation reached both regions of interest with intensities known to support phase entrainment when matched to the endogenous rhythms (28). B) During the experiment, stimulation conditions were alternated between in-phase, anti-phase and sham conditions. The 75 Hz stimulation over SMC was amplitude-modulated according to the 6 Hz stimulation over aPFC, either in-phase or anti-phase with the peaks of the 6 Hz aPFC stimulation. Sham consisted of an initial stimulation of 10 seconds, after which stimulation was terminated. C) Concurrent tACS-fMRI quantified changes in BOLD signal evoked by the tACS intervention, across in-phase and anti-phase conditions and independently from task performance. Online physiological effects of tACS are evident both under the aPFC electrode (black circle) and in other cortical regions. D) Participants with stronger inhibitory responses to theta-band stimulation over aPFC (decreased BOLD) improved their control over emotional actions (decreased congruency effect) during aPFC-SMC in-phase tACS (in green), but not during aPFC-SMC anti-phase tACS (in red), as summarized in the inset by the parameter estimates of the interaction between Emotional Control (congruent, incongruent), Stimulations Phase (in-phase, anti-phase) and tACS-Dose (BOLD signal in aPFC during stimulation vs sham (panel C); F(1,39) = 9.3, p = .004, partial eta^2^ = .19. E) Without controlling for inter-participant variability in tACS-Dose, the differential phase effect on emotional-actions control is less statistically reliable (p = .06, partial eta^2^ = .088).

We manipulated emotion-action control through a social-emotional approach-avoidance task, where human participants use a joystick to rapidly approach or avoid happy or angry faces (figure 1A). People tend to approach happy faces and avoid angry faces (*3*). Overriding these *affect-congruent* action-tendencies and instead generating *affect-incongruent* actions (approach-angry and avoid-happy) requires control, which is implemented through the aPFC, parietal/SMC and amygdala/hippocampal regions (*8, 9*). Accordingly, in the sham condition of this study, participants’ error rates and aPFC BOLD activity increased when incongruent approach-avoidance responses are compared to affect-congruent responses, reproducing previous effect sizes (*8, 9*) – Figure 1B,C.

We quantified the magnitude of the physiological effects of tACS on the underlying neural tissue through concurrent tACS-fMRI. We used BOLD effects of theta-band tACS stimulation over aPFC (in-phase + anti-phase stimulation epochs vs. sham; figure 2D), as a dose-dependent metric of tACS effects (tACS-dose (*20*)), independent of task performance. In line with our expectations, higher tACS-dose on aPFC increased the emotion-control enhancement induced by in-phase stimulation (in-phase aPFC-theta/SMC-gamma tACS) [Emotional-Control*Phase-condition*tACS-dose interaction: Figure 2E]. Interestingly, across participants, the local cortical (BOLD) effects of the inhibitory theta-band aPFC stimulation (*23*) correlated positively with improved emotion-control (indexed by decreases in behavioural congruency-effects during the in-phase condition, but not in the anti-phase condition – Figure 2D). These dose- and phase-dependent results, predicted on the knowledge that emotional-action control requires synchronization between specific endogenous rhythms (*8, 24*), indicate that in-phase aPFC-theta/SMC-gamma tACS increases participants’ control over their emotional action-tendencies.

The observed cognitive benefits of the tACS intervention could arise from direct modulation of aPFC-SMC connectivity, or be mediated by other regions (figure 3A). We arbitrated between those possibilities using two complementary approaches. First, we tested whether the amygdala mediates these effects. This region is a prime candidate as it is connected to both aPFC and SMC, and is strongly involved in emotional-action control (*9, 25*). We distinguished between a number of anatomically plausible circuit-level effects of the tACS intervention using dynamic causal modelling (*21*) on regional BOLD timeseries in amygdala, aPFC and SMC, measured during performance of the approach-avoidance task. Models comparison supports a circuit where tACS modulates aPFC→SMC, aPFC→amygdala, and amygdala→SMC connections, supplementary figure 2. Crucially, the effect of stimulation on emotional-action control was driven by tACS modulation of a specific component of that circuit, the task-related aPFC→SMC connectivity (Emotional-Control*Phase-condition*aPFC→SMC effective connectivity; Figure 3). Figure 3 visualize this behavioural- and connectivity-related difference in tACS effects, contrasting the behavioural effects evoked in participants with strong or weak in-phase tACS modulation of aPFC→SMC connectivity. Emotional control was enhanced in those participants with strong in-phase tACS modulation of aPFC-SMC connectivity: the congruency effect observed in the anti-phase and sham conditions disappeared during the in-phase condition. In contrast, emotional control remained unaffected in those participants without in-phase tACS modulation of aPFC→SMC connectivity: there were similar congruency effects across in-phase, anti-phase and sham conditions. Second, we assessed whether inter-participant variance in the circuit-level effect of the tACS intervention can be understood in terms of inter-participant variance in structural connectivity between aPFC and SMC. In support of this notion, participants with higher fractional anisotropy (a measure of white matter integrity) in the white matter beneath BA6 had stronger inhibitory coupling between aPFC and SMC during in-phase tACS (as compared to anti-phase tACS) (supplementary materials).

**Figure 3:**
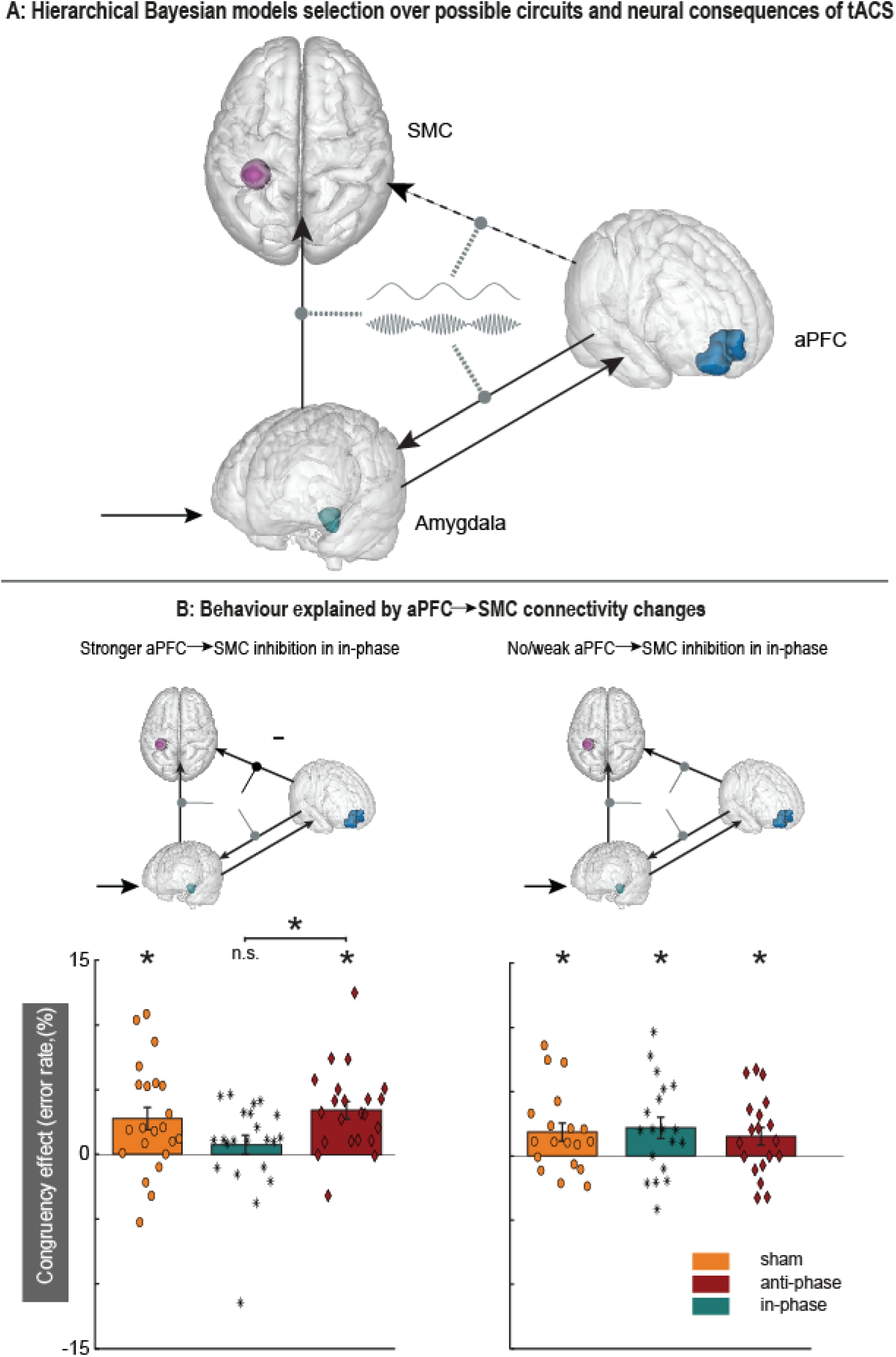
Modulatory effects of dual-site phase-coupled tACS on emotional-action control depend on effective connectivity between aPFC and SMC. A) Models selection compared models with and without a direct connection between aPFC→SMC (dashed arrow), and tACS modulations on different connections (grey dashed oval arrows). B) Model selection indicates that tACS affects multiple connections in the network (top panels; supplementary materials), but only tACS-related changes in connectivity between aPFC→SMC predict behavioural effects of the stimulation (interaction between Emotional Control (congruent, incongruent), Stimulations Phase (in-phase, anti-phase) and aPFC→SMC effective connectivity (DCM.B matrix)): F(1,39) = 6.01, p = .019, np^2^ =.13; lower panels. Those participants with stronger inhibitory influence of aPFC over SMC in the in-phase condition (lower left panel, n = 22) showed decreases in congruency effects in the in-phase condition, but not in the anti-phase and sham condition. The rest of the participant did not show a differential effect between stimulation conditions. Asterisks: p < .01.

Using a combination of concurrent tACS-fMRI, cognitively precise behavioural outcomes, as well as effective and structural connectivity, we provide converging evidence for a causal role of aPFC-SMC connections in guiding action selection during emotional-action control (*5*). Emotional-action control improves when dual-site phase-coupled tACS is tuned to increase the efficacy of anterior prefrontal inhibition over sensorimotor cortex. Alternative interpretations of the findings, focused on transcutaneous entrainment (*26*) or retinal stimulation (*27*), are ruled out by the experimental design. Namely, the in-phase and anti-phase conditions used identical stimulation parameters apart from their phase difference, thereby creating the same peripheral effects.

This study grounds a human gain-of-function intervention on the known hierarchical organization of the frontal cortex (*12*) and on phase-amplitude coupling as a mechanism for directional inter-regional neuronal communication (*12, 13*). Similar interventions could help patients suffering from social-emotional disorders. For instance, individuals with social anxiety are often unable to overcome avoidance tendencies, hampering interventions aimed to extinguish fear through exposure (*6*). Synchronizing aPFC-SMC theta-gamma coupling might temporally alleviate this lack of control (*10*), allowing patients to benefit from exposure treatment. More generally, the findings of this study pave the way for implementing physiologically-grounded non-invasive interventions aimed at reducing neural noise in cerebral circuits where communication relies on long-range phase-amplitude coupling.

## Acknowledgments

The authors would like to thank Freek Nieuwhof and Uriel PlonesThere is great interest in potentiating for their help in setting up the tACS-fMRI setup and Amy Abelmann for her assistance in data collection.

This work was supported by a VICI grant (#453-12-001) from the Netherlands Organization for Scientific Research (NWO) and a consolidator grant from the European Research Council (ERC_CoG-2017_772337) awarded to Karin Roelofs.

## Author contributions

B.B., I.T. & K.R designed research; B.B. performed research; B.B., H.d.O., I.T. & K.R. analyzed data; B.B., I.T. & K.R. wrote the first draft of the paper; B.B., H.d.O., I.T. & K.R. edited the paper.

## Conflicts declared

none.

## Supplementary materials

### Materials and Methods

#### Participants

Forty-four male students of the Radboud University Nijmegen participated in this experiment after giving informed consent. Two participants were excluded because they did not attend the whole experiment; one participant was excluded because he failed to comply with the task instructions, yielding a total pre-determined sample of forty-one participants. This sample size was determined on a priori estimates of statistical power calculated in Gpower 3.1 (*29*) according to an expected effect size of *Cohen’s d* = .4, as reported in- or calculated from (*9, 30*). The a-priori sample size calculation, and other experimental details were preregistered before onset of data analysis, and after data collection, at the Open Science Framework, (https://osf.io/m9bv7/?view_only=18ccdf99a8e84afaaebee393debbabe3). All participants had normal or corrected to normal vision and were screened for contra-indications for magnetic resonance imaging and transcranial alternating current stimulation. Mean age was 23.8 years, SD = 3.4, range 18-34.

#### Procedure

Data were acquired on three different days. On the first day, we acquired a structural T1 scan, a diffusion weighted imaging (DWI) scan, and a magnetic spectroscopy scan (MRS; not reported here). During the second and third day, electrodes were placed on the scalp, covering SMC and aPFC (see below for details on the precise localization). Afterwards, participants performed a 5 minute practice task, before starting an approach-avoidance task (35 minutes on each day) while receiving tACS stimulation and concurrently being scanned with fMRI.

#### Approach-Avoidance (AA) task

Participants performed a social-emotional approach-avoidance task that has previously been shown to require control over prepotent habitual action-tendencies to approach happy- and avoid angry faces (*3, 31*). Overriding these action-tendencies requires a complex form of cognitive control that operates on the interaction between emotional percepts and the emotional valence of the required actions (*9, 31, 32*), and depends on aPFC control over downstream regions (*5, 9, 33*), implemented by theta-gamma phase-amplitude coupling (*8*). Participants responded through a joystick with one degree of freedom (along the participant’s midsagittal plane), holding the joystick with their right hand on top of their abdomen, while laying the MR-scanner and seeing a visual projection screen through a mirror system (see below). Participants were instructed to pull the joystick towards themselves when they saw a happy face, and push it away from themselves when they saw an angry face. These were the instructions in the congruent condition. In the incongruent condition, the participants were asked to push the joystick away from themselves when they saw a happy face, and pull it towards themselves when they saw an angry face (figure 1). Written instructions were presented on the screen for a minimum of 10 seconds prior to the start of each block of 12 trials. Congruent and incongruent conditions alternated between blocks. Trials started with a fixation cross presented in the centre of the screen for 500 ms, followed by the presentation of a face for 100 ms. Participants were asked to respond as fast as possible, with a maximum response time of 2000 ms. Movements exceeding 30 % of the potential movement range of the joystick were taken as valid responses. Online feedback (“you did not move the joystick far enough”) was provided on screen if response time exceeded 2000 ms. Each participant performed 288 trials on each of the two testing days, yielding 576 trials in total, equally divided over in-phase, anti-phase, and sham stimulation – as well as over congruent and incongruent conditions.

#### tACS stimulation parameters

Transcranial alternating current stimulation (2 mA peak-to-peak) was applied using two sets of center-ring electrodes (80 mm inner Ø; 100 mm outer Ø; centre electrode had a Ø of 10 mm) (*34*). Stimulation was applied on-line during task performance in blocks of approximately 60 seconds (i.e. the length of a stimulus block; 12 trials), and consisted of a theta-band (6 Hz) sine wave over the aPFC and gamma-band (75 Hz) waveform tapered with a 6 Hz sine wave over SMC. Gamma-band power was phase locked to peaks (in-phase) or troughs (anti-phase) of the theta-band signal, figure 2B. The position of the electrodes on the skull was determined for each participant by using individual structural T1 scans to which we registered masks of the regions of interest (MNI [-28 -32 64] for SMC; (*8*); and MNI [26 54 0] for lateral Frontal Pole; (*33, 35*). Precise placement of the centre electrode was achieved using Localite TMS Navigator (https://www.localite.de/en/products/tms-navigator/; RRID:SCR_016126). Electrodes were attached using Ten20 paste (MedCat; https://medcat.ccvshop.nl/Ten20-Pasta-Topf-4-Oz,-3er-Set) and impedance was kept below 10 kOhm (mean = 3.6, SD = 2.7). Stimulation was applied using two Neuroconn DC-PLUS stimulators (https://www.neurocaregroup.com/dc_stimulator_plus.html; RRID:SCR_015520) that were placed inside a magnetically shielded box in the MR room. This box contained home-made electronics and BrainAmp ExG MR amplifiers (www.brainproducts.com) to continuously monitor the tACS output of the stimulator and filter out the RF pulses of the MR system.

#### Materials and apparatus

Magnetic resonance images were acquired using a 3T MAGNETROM Prisma MR scanner (Siemens AG, Healthcare Sector, Erlangen, Germany) using a 64-channel head coil with a hole in the top through which the electrode wires were taken out of the scanner bore.

The field of view of the functional scans acquired in the MR-sessions was aligned to a built-in brain-atlas to ensure a consistent field of view across days. Approximately 1800 functional images were continuously acquired in each scanning day using a multi-band 6 sequence, 2*2*2 mm voxel size, TR/TE=1000/34ms, Flip angle = 60°, phase angle P>>A, including 10 volumes with reversed phase encoding (A>>P) to correct image distortions.

High-resolution anatomical images were acquired with a single-shot MPRAGE sequence with an acceleration factor of 2 (GRAPPA method), a TR of 2400 ms, TE 2.13 ms. Effective voxel size was 1 × 1 × 1 mm with 176 sagittal slices, distance factor 50%, flip angle 8°, orientation A ≫ P, FoV 256 mm.

Diffusion-weighted images were acquired using echo-planar imaging with multiband acceleration factor of 3. We acquired 93 1.6 mm thick transversal slices with voxel size of 1.6 × 1.6 × 1.6 mm, phase encoding direction A >> P, FoV 211 mm, TR = 3350, TE = 71.20. 256 isotropically distributed directions were acquired using a b-value of 2500 s/mm^2^. We also acquired a volume without diffusion weighting with reverse phase encoding (P >> A).

#### Behavioural analyses

We compared differences in error rates between congruent and incongruent trials for the different stimulation conditions whilst controlling for dose dependent effects of tACS (“tACS-dose”) (*20, 36*). tACS-dose was estimated by taking BOLD contrast between stimulation and sham conditions extracted from right aPFC(*20*). This three way interaction (Emotional-Control*Phase-condition*tACS-dose) was assessed using congruency effects estimated from participant-by-participant averages (with RM-ANOVA), as well as trial-by-trial data (with Bayesian mixed effect models). RM-ANOVA was used to facilitate comparisons with earlier studies reporting on this task (e.g. (*8, 9, 32*)) and implemented in JASP (https://jasp-stats.org/; RRID:SCR_015823). We used Bayesian mixed effect models (implemented in R 3.5.3 using the brms package (*37*) because they are robust to potential violations of normality or homoscedasticity. The Bayesian mixed models included random intercepts for all subjects and random slopes for all fixed effects (congruency- and stimulation condition) and their interaction per participant. This model adheres to the maximal random effects structure (*38*). Outputs of these models are log odds with credible intervals (“*B*”). In these analyses an effect is seen as statistically significant if the credible interval does not contain zero with 95% certainty.

We hypothesized that the congruency effect in error rates would decrease for in-phase condition and increase for anti-phase stimulation and that the size of the effect per participant would depend on the BOLD effect of tACS versus sham, a measure of dose dependence that is orthogonal to the contrast of interest (in-phase versus anti-phase). These expectations were preregistered at the Open Science Framework: (https://osf.io/m9bv7/?view_only=18ccdf99a8e84afaaebee393debbabe3).

#### Modelling of stimulation currents

Current density under the electrodes was simulated using SIMNIBS version 3.1 (https://simnibs.github.io/simnibs/build/html/index.html) (*39*). We used the template head model and standard conductivities provided by SIMNIBS. Electrodes were placed over SMC and aPFC and direct currents of 1 mA were estimated to run from the inner electrode towards the outer ring. Electrode placement and current density estimates for both electrode pairs are visualized in figure 2A.

#### fMRI analyses – data preprocessing

All processing of the images was performed using MELODIC 3.00 as implemented in FSL 6.0.0 (https://fsl.fmrib.ox.ac.uk). Images were motion corrected using MCFLIRT (*40*), and distortions in the magnetic field were corrected using TOPUP (*41*). Functional images were rigid-body registered to the brain extracted structural image using FLIRT. Registration to MNI 2 mm standard space was done using the nonlinear registration tool FNIRT. Images were spatially smoothed using a Gaussian 5 mm kernel and high pass filtered with a cut-off that was automatically estimated based on the task structure. Independent component analysis was run with a pre-specified maximum of 100 components (*42*); these components were manually inspected to remove potential sources of noise (*43*).

#### fMRI analyses – signal-to-noise (SNR) artefacts resulting from electrode presence

To assess whether the presence of electrodes on the scalp had an effect on signal to noise ratio in the fMRI signal we estimated temporal signal to noise ratio (tSNR) from right aPFC (electrode present) and left aPFC (electrode not present). tSNR was calculated by dividing the mean of the signal over time by the standard deviation, separately for left and right aPFC. These estimates were extracted from a mask of the lateral Frontal Pole (*35*). We also compared the mean signal extracted from right-with left aPFC.

#### fMRI analyses - GLM

First and second level GLM analyses were performed using FEAT 6.00 implemented in FSL 6.0.0. The first-level model consisted of twelve task regressors: Approach angry, approach happy, avoid angry and avoid happy trials were modelled separately for each stimulation condition (in-phase, anti-phase and sham). In each regressor, each event covered the time interval from presentation of a face until the corresponding onset of the joystick movement. Estimated head translations/rotations during scanning (six regressors), temporal derivatives of those translations/rotations (six regressors), and MR-signals in white matter and cerebrospinal fluid (2 regressors) were included to the GLM as nuisance covariates. We considered the following comparisons. Emotional control effects were estimated by comparing incongruent trials (approach angry and avoid happy) to congruent trials across the three stimulation conditions (total congruency effect), as well as separately for each stimulation condition (in-phase, out-of-phase, and sham congruency effects). Overall stimulation effects were estimated by comparing in-phase and anti-phase stimulation conditions to the sham condition, aggregated across congruent and incongruent conditions, yielding a “tACS-dose” measure. Phase-dependent stimulation effects were estimated by comparing in-phase stimulation to anti-phase stimulation, across congruent and incongruent conditions. First level models of the two separate sessions were combined using Fixed Effects analyses implemented in FEAT. Group effects were assessed using FLAME 1 with outlier de-weighting (*44*), making family-wise error corrected cluster-level inferences using a cluster-forming threshold of z > 2.3. This threshold provides a false error rate of around 5% when using FSL’s FLAME 1 (*45*).

#### fMRI analyses – Dynamic Causal Modelling

Dynamic causal modelling (DCM 12.5), implemented in SPM 12, was used to make inferences on network effects of the tACS manipulation (*21*). Dynamic causal modelling is an approach that aims to infer hidden neuronal dynamics from neuroimaging data (*21, 46*). It requires the specification of plausible generative models describing how neural activity leads to observed neuroimaging data through haemodynamic functions. This approach allows formal comparison on different models explaining the same data (*47, 48*). DCM provides estimates of effective connectivity between neural populations, and its modulation by experiment conditions (e.g. in this case tACS present or absent). We pre-registered the hypothesis that changes in connectivity between aPFC and SMC due to tACS might be mediated by the amygdala, a region linked to regulation of social-emotional action tendencies (*9, 33, 49*). To test this hypothesis, we constructed a model space in which each model contained three regions; left Amygdala (see GLM results); right aPFC; and left SMC.

The regions of interest were defined separately for each participant based on their functional effects (from the GLM analyses), but constrained to a-priori determined regions; right aPFC, based on the FPl mask (*7, 12*); left SMC, based on a previous study involving the same task [MNI: -28 -32 64] (*8*); and left Amygdala, based on the Automated Anatomical Labelling mask (*9, 49, 50*). The timeseries of the first eigenvariate across all significant voxels in the ROI for amygdala and SMC were extracted from a sphere with 3 mm radius (amygdala) and 5 mm radius (SMC) around the peak voxel detected in each ROI in a GLM *F*-contrast across effects of interest. aPFC time series were extracted from a sphere with 3 mm radius around the peak voxel in the incongruent > congruent t-contrast in right lateral Frontal Pole (*35*).

Model structure for all models under comparison was based on the minimal architecture needed to dissociate whether synchronisation between aPFC and SMC was gated by the amygdala or via other region(s). All models consisted of a directed connection between amygdala and SMC (*51*) (indicated as amygdala→SMC), amygdala → aPFC, and aPFC → amygdala (*33, 52*). A subset of models also included aPFC → SMC (figure 3), which captures connectivity between aPFC and SMC gated via cerebral structures other than the amygdala given that the two structures do not share a monosynaptic connection (*35*). Return connections SMC → amygdala and SMC → aPFC were not included. We assumed these to be unnecessary to explain effects of synchronisation given the hierarchical nature of prefrontal control (*53, 54*). The onset of the stimulus in each trial (presentation of the face) was taken as a driving input (DCM.C matrix) feeding into the amygdala node of each model. Emotional control (all incongruent trials) and tACS condition (all in-phase trials, all anti-phase trials or a combination; Supplementary Figure 3B) were allowed to modulate different connections (DCM.B matrix).

We created 84 models that systematically varied in four dimensions of interest, which were compared using family-wise model comparison (*48*). Dimension 1 was the modulation of emotional control, where control was allowed to modulate either amygdala → aPFC connection or aPFC self-connections (*32*). The second dimension of interest was the connectivity structure of the models. This dimension consisted of two different model types: with or without a direct connection from aPFC → SMC (Sup. Figure 2A). Presence of this connection can account for influence of aPFC on SMC that is not gated by the amygdala. Dimension 3 reflects the nature of the tACS effects (Supplementary figure 2B; (*55*)). We modeled no effect of tACS (Stim 0); only the in-phase condition (Stim 1); only the anti-phase condition (Stim 2); both conditions separately (Stim 3); both conditions with opposite sign and similar amplitude (Stim 4). Dimension 4 arbitrated over the location of tACS modulation (figure 3A, Supplementary figure 2C), which was allowed to modulate amygdala → SMC, aPFC → amygdala, and aPFC → SMC, or any combination of the three connections. We differentiated these possibilities by separating all models into model families for each dimension, allowing inferences per dimension while averaging over all other parameters (Sup. Figure 2) (*48*).

After model comparison, we extracted connectivity estimates that were altered by tACS from the winning model and used those parameters to predict tACS effects on behaviour. The estimated changes in connectivity due to tACS are estimated independently of the behavioural effects of tACS, and were used to test for an interaction between Emotional Control (congruent, incongruent), Stimulations Phase (in-phase, anti-phase) and effective connectivity (DCM.B matrix) using RM-ANOVA and Bayesian mixed effects models.

#### DWI analyses

All analyses of diffusion data were performed in FSL’s FDT 3.0 (https://fsl.fmrib.ox.ac.uk). We used TOPUP to estimate susceptibility artifacts using additional b = 0 volumes with reverse phase coding direction (*41*). Next, EDDY correction was used (using the fieldmap estimated by TOPUP) to correct distortions caused by eddy currents and movement (*56*). We used BedpostX to fit a crossing fiber model with three fiber directions (*22*). Connections between aPFC and BA6 were reconstructed using *probtrackx2.* Seed mask was the lateral Frontal Pole (*35*), which is a mask that contains voxels bordering white matter. Target mask was area BA6 contained within the Juelich atlas in FSL. This probabilistic mask was thresholded to contain only voxels that were at least 50% likely to be in BA6. Tractography was run twice with the standard settings recommended in probtrackX and an exclusion mask in the midline. Estimated connection strengths were normalized to unit length within participants. We averaged all connections over participants and thresholded this volume at .5 to create a mask of likely pathways linking FPl (within aPFC) and BA6. In a second step we constrained the tractography to either only include connections via the thalamus, or only via medial and lateral pathways through the prefrontal cortex.

Tract based spatial statistics (TBSS (*57*)) was used to assess whether the white matter integrity in the voxels in this connection mask explained part of the variance in response to tACS. TBSS was run using the default settings provided by FSL. In short, FA images were eroded and nonlinearly registered to standard space. After this we derived a mean skeleton based on all participants and thresholded the skeletonized FA values at .2. We then used *randomise* (*58*) with threshold free cluster enhancement (*59*) to make inferences on the correlation between FA integrity and DCM effective connectivity estimates (in-phase versus anti-phase). To reduce the search space, the comparison was constrained to voxels that were part of the mask linking FPl to BA6 with and without the constrained connectivity (see supplementary figure 2).

## Supplementary Results

Participants performed a social-emotional approach-avoidance task in which they approached or avoided happy and angry faces using a joystick. Approaching angry- and avoiding happy faces is incongruent with automatic action tendencies to approach appetitive and avoid threatening stimuli, and requires control over these prepotent habitual action tendencies (figure 1; (*9, 31, 33*). fMRI was acquired during task performance whilst participants received on-line dual-site theta-gamma tACS over aPFC and SMC, figure 2A;B. Control over social-emotional action tendencies involves phase-amplitude coupling between prefrontal theta- and SMC gamma-band rhythmic activity (*8*), the phase organisation of which is best reflected by the in-phase condition. In-phase, anti-phase and sham conditions (figure 2B), were alternated throughout the task, allowing comparison of on-line, within-subject effects of stimulation conditions.

### Signal quality below the electrodes

There was a reduction in absolute signal in right aPFC below the frontal electrode as compared to the contralateral aPFC, t(40) = 5.5, p <.001. This consisted of an average reduction in signal intensity of 11% (SD 10%). However, there was no difference in temporal SNR in right aPFC as compared to left; t(40) = .2, p = .7. These results are in line with earlier reports on noise induced by the presence of electrodes and stimulation equipment in the scanner bore (*60*). Although the absolute MR signal is reduced due to the presence of the tACS electrode, there is no effect on the SNR of the underlying brain region, suggesting that condition differences can be estimated reliably. This is supported by the consistency between the results obtained in the congruency contrast across stimulation conditions, and the results obtained in earlier reports using the same paradigm (figure 1C, supplementary figure 1; (*9, 31, 33*))

### Behavioural and neural costs of controlling social-emotional action tendencies

Across all conditions (stimulation and sham combined), reaction times were longer in the incongruent (M = 687 ms, SD = 141) than in the congruent condition (M = 637 ms, SD = 111), brms estimate *B* = 23.1 ms, credible interval *(CI) [15 31]*; paired t-test congruent vs incongruent *t*(40) = 5.9, *p* < .*001*, Cohen’s d *(M1-M2/SD*_*pooled*_) = .39 [.22 1]. Participants also made more errors in the incongruent condition (94.8 % correct, SD = 3.4 %) than in the congruent (96.9% correct, SD = 2.3 %), this was significant for both aggregated scores: *t*(40) = 5.4, *p* < .001, *d* = .7 [.01 1.35], as well as on the trial-level: *B = 0.29 CI [.16 .42]*. The latter parameter is a log odds and indicates that participants are more likely to make correct responses in the congruent than incongruent trials. These effects illustrate the behavioural cost of controlling automatic action tendencies.

Whole-brain cluster-corrected fMRI analyses show that controlling social-emotional action tendencies elicited stronger BOLD activity in bilateral prefrontal cortex: MNI [-18 54 -16] and [40 42 32], extending into lateral Frontal Pole (*33, 35*); bilateral posterior parietal cortex (PPC) MNI: [38 -58 38] and [-56 -44 46]; precuneus [6 -62 72], temporal cortex [-66 -48 0]; and inferior frontal gyrus [60 20 4], figure 1C, supplementary figure 1C. Activity in the parahippocampal gyrus [-24 -40 -14], extending into hippocampus and amygdala; medial prefrontal cortex [0 66 2]; and lateral occipital cortex [52 -74 -10] was stronger for congruent than for incongruent trials. Similar effects have been reported in several earlier reports (*9, 33, 49*), with prefrontal effects depending crucially on the interaction between the emotional content of the action and the emotional content of the percept, over and above the emotional value of the action or stimulus alone (*9, 31, 61*).

### Cerebral effects of dual-site phase-coupled tACS versus Sham

During task performance each participant received concurrent transcranial alternating current stimulation on two locations; aPFC (*33, 35*) and sensorimotor cortex, figure 2A;B. These locations, frequency, and phase relationship were based directly on a previous study showing aPFC-SMC phase-amplitude coupling in theta-gamma band rhythmic signals (*8*). Whole-brain cluster-corrected analyses contrasting both stimulation conditions versus sham showed increased activity in several brain regions, most of which overlapped with task relevant regions: right prefrontal cortex [42 16 48], extending into FPl [22 60 0]; bilateral posterior parietal cortex [50 -58 50] and [-56 -44 46], figure 2C, supplementary figure 1C;D. These effects support previous experimental and theoretical work suggesting that tACS effects on-going neural activity and effects are task-dependent (*62*).

### Cerebral effects of in-phase vs anti-phase dual-site phase-coupled tACS

Contrasting in-phase with anti-phase conditions showed stronger activity in left aPFC [-28 50 - 10], contralateral to stimulated right aPFC, and left PPC [-56 -46 52], supplementary figure 2E. These results suggest that desynchronising right aPFC and left SMC might induce compensatory effects in the contralateral homotopic control regions. These findings illustrate the importance of measuring online effects of brain stimulation because stimulation might have distal effects on neural tissue (*63*).

### Dose dependent effects of tACS on social-emotional control

To assess whether the tACS intervention increases control over social-emotional behaviour we compared behavioural congruency effects between different stimulation conditions whilst controlling for dose dependent effects. Dose dependent responses were calculated by extracting the differential BOLD signal under the prefrontal stimulated region (right aPFC, [22 60 0]) when comparing stimulation versus sham epochs. This BOLD effect provides a participant-by-participant index of aPFC response to the tACS intervention that is orthogonal to the in-phase versus anti-phase comparison.

Following the expectation preregistered at the Open Science Framework: (https://osf.io/m9bv7/?view_only=18ccdf99a8e84afaaebee393debbabe3), there was a three way interaction between Emotional-Control, Phase-condition, and tACS-dose, both on the trial level; *B = .1 CI [.007 .2]*, and in aggregated error rates; RM-ANOVA, *F(1,39)* = 9.3, *p* = .*004*, partial eta^2^ = .19, indicating that participants performed differently in the two stimulation conditions as a function of the strength of the tACS influence. Post-hoc correlation analyses between BOLD change due to tACS and congruency effects showed that these effects are mainly driven by the in-phase condition: Spearman’s Rho = .32, p = .04, figure 2B. This finding indicates that, in the in-phase condition, larger BOLD responses to the inhibitory theta-band rhythm corresponded to smaller behavioural congruency effects. In the anti-phase condition, there was no correlation between BOLD response and behavioural congruency effects: Spearman’s Rho = -.23, *p* = .14. Controlling for baseline congruency effects (in the sham condition) did not change these results; Rho = .32, p = .044 for in-phase; Rho = -.26, p = .09 for anti-phase. These results suggest that theta-gamma phase-coupled tACS increases control over social-emotional behaviour in participants that respond to stimulation. Contrasting error-rate congruency effects (% correct for congruent - % correct for incongruent trials) without accounting for the dose dependent responses did not show a statistically reliable group effect of stimulation condition on the trial level, *B* = .05 CI [-.05 .14], nor on the aggregated scores RM-ANOVA; *F(1,39)* = 3.75, *p* = .*06*, partial eta^2^ = .088, Sup figure 1A.

Reaction times did not show differential effects in the different stimulation conditions, Emotional-Control*Phase-condition*tACS-dose; *F*(1,40) = .05, *p* = .8, congruency*stimulation; *F*(1,41) = 2.3, *p* = .14, Sup figure 1B. The lack of effects on reaction times is in line with expectations preregistered at the Open Science Framework: (https://osf.io/m9bv7/?view_only=18ccdf99a8e84afaaebee393debbabe3) as well as earlier stimulation studies using this same task (*9*). This finding suggests that the aPFC control required during the AA task operates at the level of action selection.

### tACS changes aPFC-SMC effective connectivity, which correlates with behavioural effects

To assess the network effects of dual-site tACS we used dynamic causal modeling (DCM; (*21*)) verifying whether aPFC→SMC connectivity was gated by the amygdala. We created a set of 84 models with two different connectivity structures; with and without direct connection between aPFC and SMC, supplementary figure 2A. These models differed in the way tACS could affect connectivity, Sup figure 2B. Possibilities consisted of no effect of stimulation (Stim 0), effects of one stimulation condition (stim 1 and stim 2, Sup figure 2B), independent effects of both conditions (Stim 3) or opposite sign but equal amplitude effects of both stimulation conditions (Stim 4). Finally, models varied with respect to the location of the tACS modulation: Amygdala → SMC, aPFC → amygdala or aPFC → SMC, two of those three connections, or all three connections, supplementary figure 2C.

Family comparisons showed that the structural model containing a direct connection from aPFC →SMC was more likely than a model without this connection, supplementary figure 2A. This suggests that not all effects of aPFC→ SMC are mediated by the amygdala. Comparing the different possible effects of stimulation showed that models in which both stimulation conditions were modeled separately had the strongest model evidence, figure 2B. Finally, comparisons over possible locations where tACS could modulate connectivity showed tACS effects on all three modeled connections, supplementary figure 2C

We extracted connectivity parameters from all three modulated connections and used them to account for differences in behavioral response. This showed that differences in effective connectivity between in- and anti-phase on the aPFC → SMC connection predicted differences in behavioral effects between stimulation conditions both on the trial level; *B = .11 CI [.02 .2]* and on aggregated congruency-effects; RM-ANOVA; *F(1,39)* = 6.01 *p* = .019, np^2^ =.13. Over the whole group connectivity parameters did not differ between in- and anti-phase conditions, *t*(*40*), = *1*.*1, p* = .*27*. Post-hoc tests showed that those participants that showed more negative connectivity between aPFC→SMC in the in-phase as compared to anti-phase condition showed a reduction in congruency effects in the in-phase as compared to the anti-phase condition, t(21) = -2.9, *p* = .*008*. This was not the case for participants with no difference, or more negative connectivity for anti-phase versus in-phase, *p* = .*35*; main figure 3B. This improvement of control in the in-phase condition remains when outliers in behavioral congruency are removed, *t*(*20*), = - *2*.*7, p* = .*012*, and if we remove the participants that did not show a response to amygdala input in the DCM analysis (n = 3), *t(19)* = -*2*.*8, p* = .*01*. Strikingly, for the participants with stronger negative connectivity between aPFC→SMC the congruency effect was no longer significantly different from zero, *t(21)*, = .99 *p* = .33 in the in-phase condition, indicating that they did no longer perform worse when having to override automatic action-tendencies. Participants that did not show tACS modulation of aPFC→SMC connections in the in-phase condition did show a congruency effect *t(18)* = 2.6, *p* = .*018*. These effects suggest that successful synchronization of aPFC-SMC rhythmic activity increases control over social-emotional behavior by increasing aPFC inhibition over SMC.

To explore whether structural characteristics might underlie individual differences in response to stimulation we tested whether differences between aPFC→SMC connectivity parameters between in-phase and anti-phase conditions correlated with FA integrity in voxels potentially connecting aPFC to SMC. We observed a correlation between FA integrity and DCM connectivity estimates for in-phase versus anti-phase in white matter regions leading up to right BA6, *tfce* corrected for voxels in a thresholded mask connecting FPl (aPFC) and BA6; *p* = .*046*, supplementary figure 3. Here, increased inhibition in the in-phase condition (relative to anti-phase) is related to higher FA integrity in white matter leading up to BA6. Further exploration showed that this correlation is present when considering cortico-thalamo-cortical connections; corrected *p* < .*048*, but not when only considering cortico-cortical connections. These findings suggest that tACS synchronization of aPFC→SMC connectivity might partly depend on structural integrity in aPFC-thalamic-SMC connectivity (*64*). However, given that these results are exploratory and depend on the threshold chosen for creating the masks they need to be considered with caution and warrant replication.

---

**S Figure 1:**
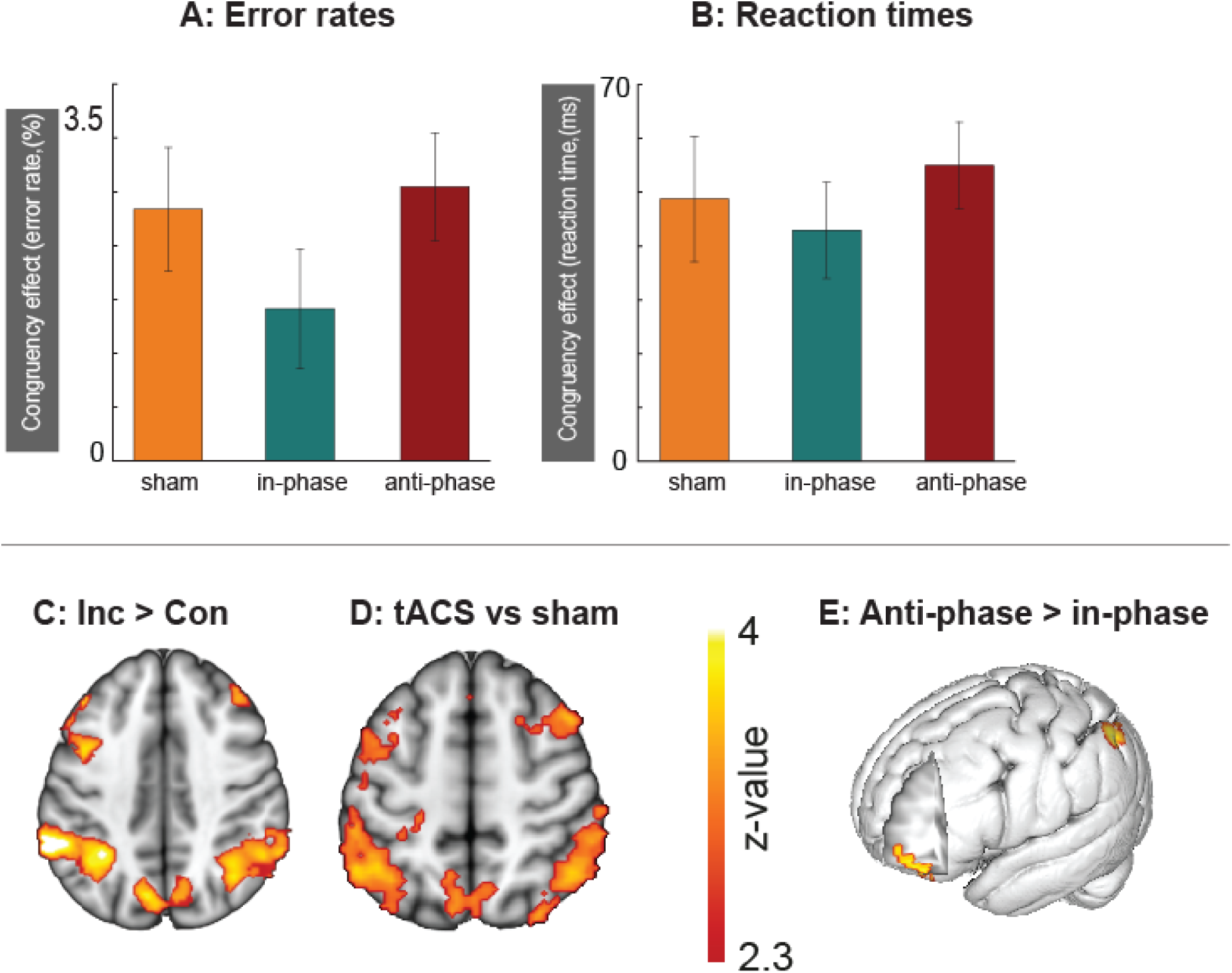
Behavioral and neural effects in different task conditions. A) Average congruency-effects in percentage correct over all three conditions. Differences between in-phase and anti-phase were trend-significant (p = .06) when not controlling for dose dependence. B) Average reaction time congruency effects, visual inspection shows similar patterns as the error-rates but there are no statistical differences between conditions. Reaction time effects were not anticipated based on earlier brain-stimulation studies using this task (9). C) Increased activity in posterior parietal and premotor areas for incongruent versus congruent and D) stimulation versus sham. E) Contrasting in-phase with the anti-phase condition shows increased activity in left aPFC and PPC, contralateral to the prefrontal stimulation site. Speculatively, these effects might suggest network compensatory effects when right aPFC is desynchronized from the control network.

**S Figure 2:**
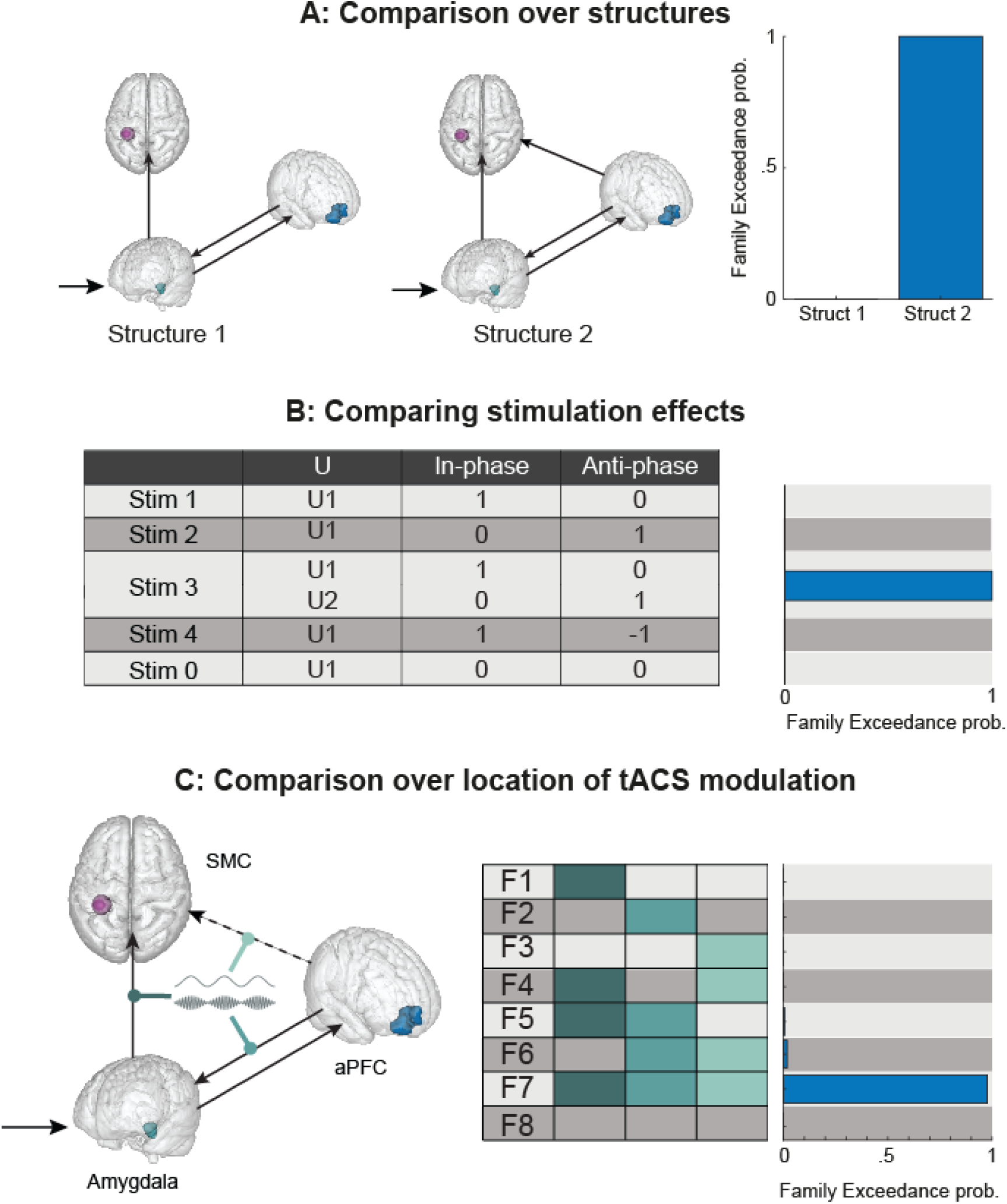
Comparing models of tACS modulation. A) Comparison over structure families. Structure 1 does not contain a direct connection between aPFC and SMC, tACS modulation of aPFC→ SMC connectivity is therefore forced through the amygdala. Structure 2 does contain a direct connection. Model comparison shows that structure 2 is more likely, suggesting that not all modulation of aPFC→ SMC is gated by the amygdala. B) Comparison over stimulation effects. There were five ways in which the tACS manipulation could modulate connectivity in our models. “Stim 1” and 2 model effects of either in-phase or anti-phase effects respectively. “Stim 3” models in- and anti-phase effects to modulate connectivity independently of one another. “Stim 4” models in-phase and anti-phase as being opposite in sign but similar in amplitude. “Stim 5” models no effect of stimulation in either condition. The data is best described by Stim 3, showing independent effects of both stimulation conditions on connectivity. C) Comparison over stimulation locations. tACS was allowed to modulate three different connections; i) connection from aPFC → amygdala, ii) connection from amygdala → SMC and iii) connection from aPFC → SMC. We also included models having two, or all connections modulated by tACS. Model comparison shows that all three connections are influenced by tACS, suggesting network wide effects of stimulation. Only parameters extracted from aPFC→SMC connections predicted behavioral effects.

**S Figure 3:**
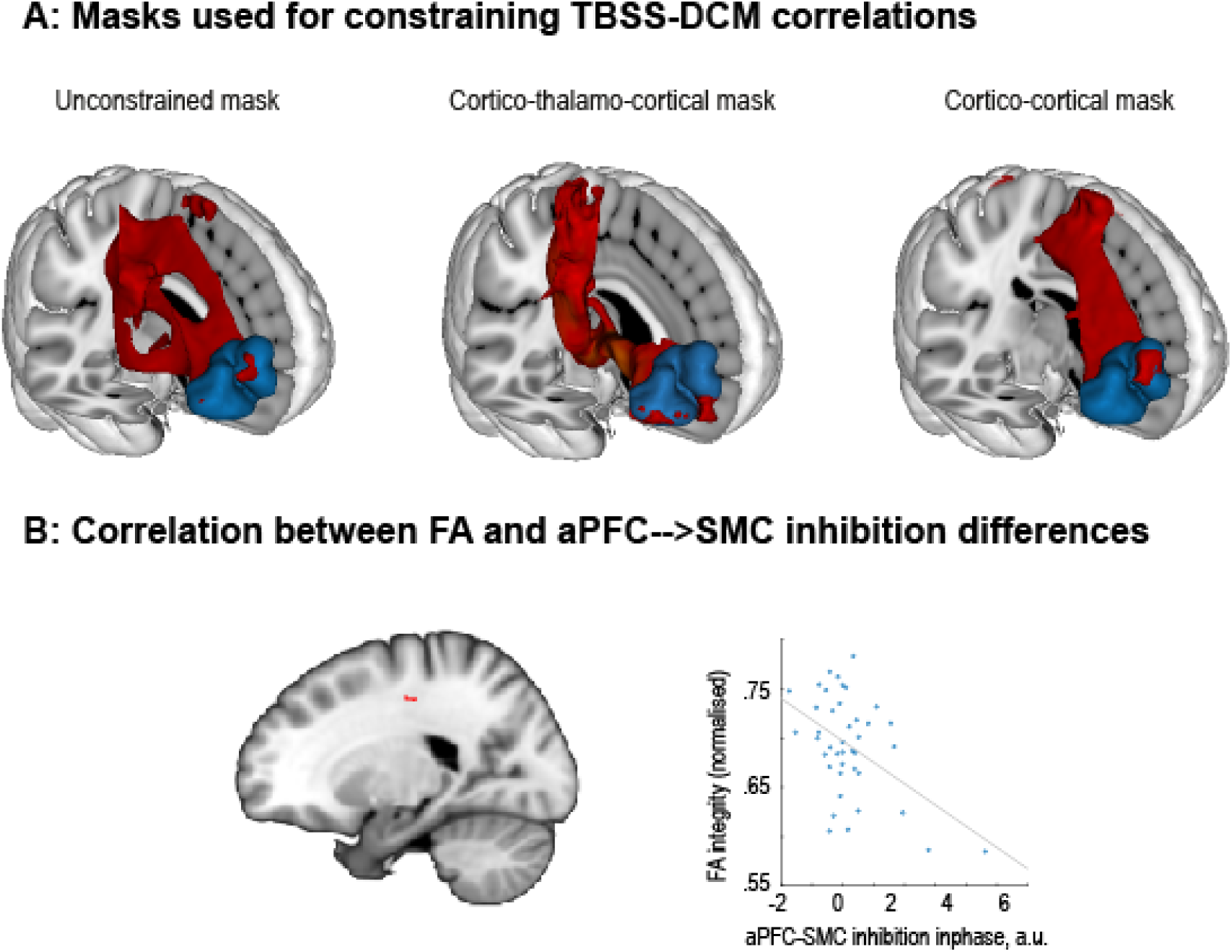
Exploratory correlations between tACS modulation and FA values. A) Masks (in red) that are used as region of interest for correlating aPFC→SMC connection estimates from DCM with FA values. The first mask was created by unconstrained probabilistic tractography between FPl (aPFC, in blue) and BA6. The middle mask was constrained to go through the thalamus. The third mask was created by constraining tractography to cortico-cortical connections. B) FA values in voxels contained in tractography masks connecting FPl (aPFC) and BA6 via the thalamus (middle mask in A) correlated with increased inhibition of aPFC→SMC in the in-phase versus anti-phase. Here people with stronger FA values showed more inhibition in in-phase versus anti-phase.

